# Sympatric wren-warblers partition acoustic signal space and song perch height

**DOI:** 10.1101/754606

**Authors:** Shivam S. Chitnis, Samyuktha Rajan, Anand Krishnan

## Abstract

By evolving divergent acoustic signals, sympatric assemblages of animals may minimize potentially costly masking interference. Acoustic signal space may be multidimensional, with coexisting species also vocalizing from different regions of physical space. Here, we demonstrate acoustic signal space partitioning in four sympatric species of wren-warbler (Cisticolidae, *Prinia*), in an Indian dry deciduous scrub habitat. We find that the breeding songs of wren-warblers are divergent from each other in multivariate parameter space, with only minimal interspecific overlap. Partitioning of signal space exerts constraints on the intraspecific diversity of acoustic signals, and each species exhibits different strategies to overcome these constraints. Two species partition intraspecific signal space into multiple note types, whereas a third exhibits intraspecific variation in repetition rate, thus supporting song diversity within a constrained acoustic space. Finally, we find that the four species also partition song perch heights, thus exhibiting separation along multiple axes of acoustic signal space. We hypothesize that divergent song perch heights may be driven by competition for higher singing perches or other ecological factors rather than signal propagation. Acoustic signal partitioning along multiple axes may therefore, we propose, arise from a combination of diverse ecological processes.

## Introduction

The diversity of animal acoustic signals is thought to have evolved under the competing pressures of remaining conspicuous to conspecifics, and simultaneously minimizing the risk of predation or erroneous recognition(Wilkins et al. 2013). The latter is particularly relevant in sympatric groups of closely related species, where errors in species recognition are costly(Grant and Grant 2010). Considerable research has therefore focused on how the signals of closely-related species diverge(Henry and Wells 2010). In multispecies assemblages, interspecific divergence may result in a pattern where each species occupies a “niche” in signal space that is distinct from all others(Chek et al. 2003; Schmidt et al. 2013). For birds, which use acoustic signals to defend territories and attract mates, divergent acoustic signals enable them to communicate with minimal competitive or masking interference from heterospecifics(Luther 2009). Signal space partitioning may occur along multiple axes; competition for singing perches(Krams 2001; Sprau et al. 2012), or constraints related to maximizing sound transmission(Marten and Marler 1977; Nemeth et al. 2002; Barker and Mennill 2009) may result in sympatric species also partitioning physical space (i.e. singing perches) within the environment. This may further minimize acoustic overlap between them.

Partitioning acoustic signal space along multiple axes, including evolving divergent frequency time parameters and singing from different locations, imposes constraints on the intraspecific signal space of each species. By evolving diverse note types, or varying the temporal pattern of songs, individual species may further partition intraspecific signal space, thus employing complex communication in spite of the constraints of the “acoustic niche”. Signal space partitioning has been studied extensively in frogs and crickets (Duellman and Pyles 1983; Chek et al. 2003; Schmidt et al. 2013). In birds, which often possess highly diverse and complex songs, frequency and time-partitioning may help minimize acoustic signal interference(Fleischer et al. 1985; Brumm 2006; Luther 2008; Krishnan and Tamma 2016). Additionally, a few studies have found perch height segregation in forest birds, attributed to maximizing sound propagation(Nemeth et al. 2002; Barker and Mennill 2009; Tobias et al. 2010), minimizing predation risk or male-male competition(Krams 2001; Sprau et al. 2012). However, signal space partitioning in birds along multiple axes remains relatively understudied. Because of the varied note types in the learned songs of passerine birds, studying sympatric groups of related species may help understand how they overcome the limitations of their signaling niche. Additionally, sound propagates differently in more open scrub/grassland habitats(Boncoraglio and Saino 2007), which are still underrepresented in tropical bioacoustics studies, and it is thus important to study the acoustic signal space of birds occupying these habitats as a comparison to forest birds.

Here, we study signal space partitioning along the axes of signal parameter space and physical singing space (perch height) in four sympatric wren-warblers (Cisticolidae, genus *Prinia*) occupying a seasonally dry scrub-grassland habitat in Maharashtra, Western Peninsular India. These four congeners breed during the Southwest monsoons, during which they utter conspicuous breeding songs. All four species are some of the most abundant singing birds in this habitat(Krishnan 2019), suggesting that they are likely to encounter significant interspecific overlap across all axes of acoustic signal space. We utilized a focal recording approach to describe inter- and intraspecific variation in *Prinia* acoustic signals, and also recorded the perch heights at which birds sang their songs. Our study illustrates the diversity of strategies that may be employed by closely related birds to minimize masking interference and competition for signaling space.

## Materials and Methods

### Study site and species

We conducted this study at the Vetal Tekdi Biodiversity Park, a small remnant of dry *Acacia-Anogeissus* scrub-grassland mosaic with scattered trees in Pune, Maharashtra, India(Nerlekar and Kulkarni 2015). Four sympatric species of wren-warbler or prinia (genus *Prinia*) occur within this landscape, the Ashy (*P. socialis*), the Grey-breasted (*P. hodgsonii*), the Plain (*P. inornata*), and the Jungle (*P.sylvatica*) prinias (Figure 1). Our study was timed to coincide with the onset of the breeding season, with recordings conducted in June and July 2018 during the early monsoon season.

**Figure 1:**
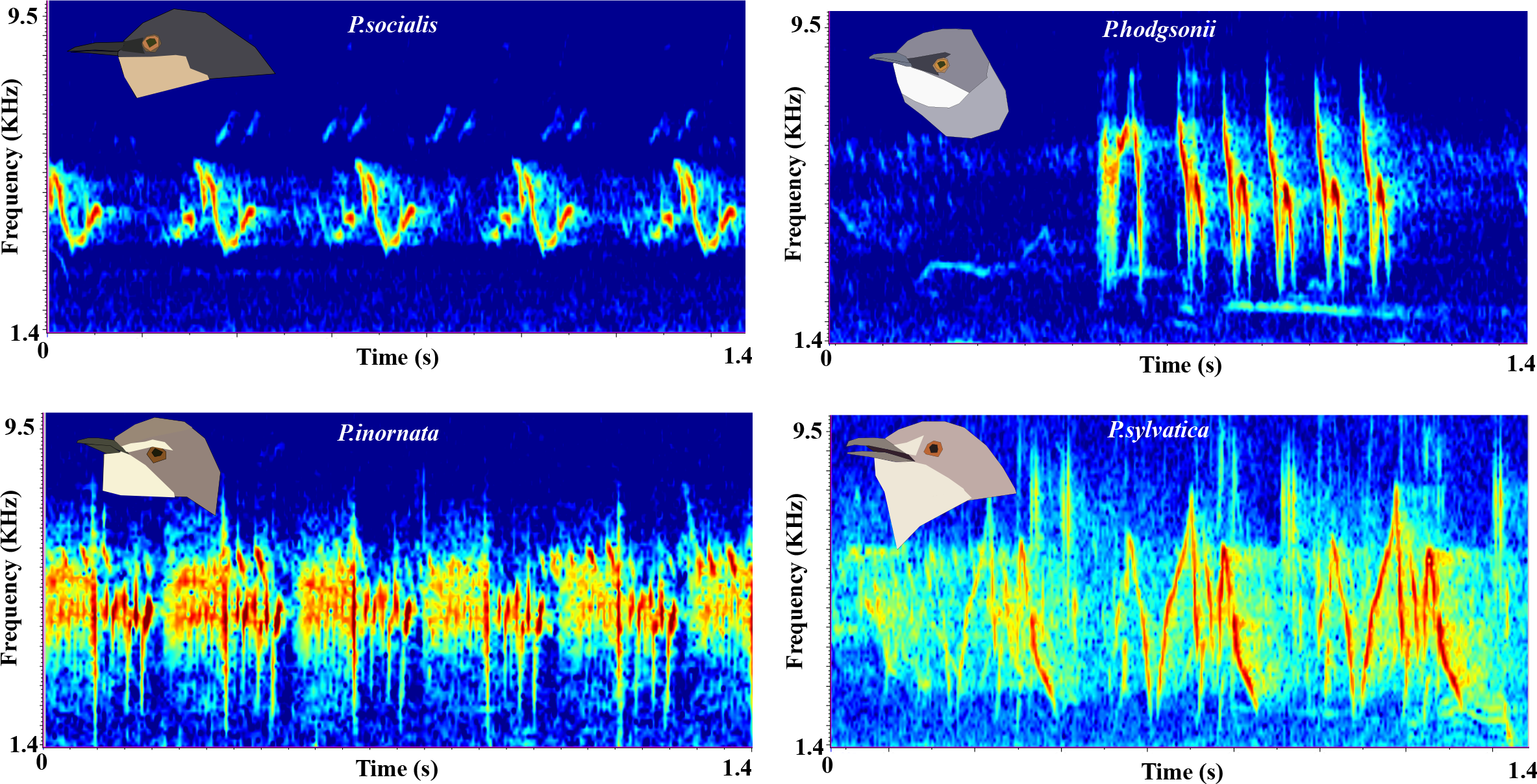
Breeding song spectrograms of four sympatric *Prinia* wren-warblers from Pune, Maharashtra.

### Recording

Our recordings were made during the morning peak in singing activity, between 6-10AM. First, we located singing birds and recorded their vocalizations using a Sennheiser ME62 (Wedemark, Germany) omnidirectional microphone connected to a Zoom H6 (Tokyo, Japan) handheld recorder. Each recording began as soon as the bird was located, and continued until the bird left the singing perch. We kept notes on each recording both in a notebook and verbally at the end of each recording, and transcribed them into datasheets after sampling sessions were concluded. After concluding a single recording, we estimated the song perch height (see below), made notes on species, type of vocalization (calls versus song) to match sound files to measurements, and then proceeded to locate another vocal bird. Our total sampling effort at the end of data collection resulted in about 160 individual measurements totaling approximately 4.5-5 hours of audio, from which we measured a total of 4615 song notes across species (*P.socialis* N=1799, *P.hodgsonii* N=1356, *P.inornata* N=652 and *P.sylvatica* N=806).

### Analyses

Using Raven Pro 1.5 (Cornell Laboratory of Ornithology, Ithaca, NY, USA), we labelled song and call notes of all four species (Hann window size 512 samples with overlap of 256 samples) and calculated the following nine parameters for each note: note duration, 90% bandwidth, center frequency, average entropy, average peak frequency, maximum and minimum peak frequency, and the peak frequency at the start and end of the note (calculated using the peak frequency contour feature in Raven Pro). We were careful to exclude recordings with high background noise to avoid confounding variability in measured parameters. To visualize the signal parameter space occupied by *Prinia* acoustic signals, we performed a principal components analysis on the correlation matrix of all parameters measured (combining call and song notes, as defined in the literature)(Rasmussen and Anderton 2005), and ordinated the points corresponding to song notes in principal component space. Further, to examine how distinct the song notes of different species were from each other, we carried out a linear discriminant analysis (LDA) on the note parameters of the breeding songs, using the Classification Learner app in MATLAB (Mathworks Inc., Natick, MA, USA). This analysis trained a linear discriminant model and assessed its accuracy at classifying notes to species using 10-fold cross-validation. We additionally performed a separate LDA on call notes (452 total notes across species) as well, to examine interspecific differences between them. To further statistically test for partitioning of note parameter space (considering notes of the breeding songs only), we additionally performed two types of randomization analysis to quantify whether average interspecific Euclidean distance in PC space was greater than expected by random chance (indicating less overlap). In the first, we generated random distributions for each PC (Supplementary Figure 1) spanning the same range of values, and then sub-sampled each PC at random to generate a 4-species “null community”, where each species had the same sample size as the four in our dataset. Additionally, we performed a second randomization by randomly reshuffling data points across species within our existing dataset to obtain a second type of “null community”. For both methods, we then calculated the average interspecific distance for each of 10000 such randomized “null communities”, and measured the Z score of the observed average interspecific distance with respect to this randomized distribution. We predicted that a partitioned song space would lead to a greater interspecific distance than random chance, and therefore a significantly positive Z score. Finally, we performed randomizations using the first method for each species pair (6 in total), to examine whether each pair exhibited greater interspecific distance than expected by random chance. This would indicate that each species occupied a unique region of multivariate signal parameter space.

To examine whether species possessed multiple song note types within intraspecific song signal space, we first examined the notes visually to identify broad groupings within species. Next, we performed hierarchical clustering using the cluster function in MATLAB to empirically verify whether clusters obtained using the previous method were indeed well-separated. As a final quantification of the accuracy of this method, we performed a third LDA on these clusters to assess how distinct and well-separated they were from each other. For species with only a single note type, we determined whether they instead varied the repetition rate of their signals, by examining variation in the interval between notes (the time between the end of one note and the beginning of the next).

### Perch height measurements

We estimated song perch heights in the field after each focal recording. Because of the inherent uncertainty in estimates of perch height taken in the field, we grouped song perches into four broad categories: perches <2m (grass and lower branches of bushes), 2-5m (tops of bushes), 5-10m (small trees and lower branches of large trees), and >10m (tops of large trees). To statistically test whether perch height distributions across species differed from chance, we used a χ^2^ test against an expected distribution assuming no interspecific patterns in perch height.

## Results

### Prinia species exhibit partitioning in acoustic signal parameters

The first three principal components of nine acoustic parameters explained over 77% of the variation in *Prinia* acoustic signals (both songs and calls) (Table 1). PC1 loaded moderately positively on all frequency parameters and moderately negatively on bandwidth and note duration, whereas PC2 loaded moderately to strongly on entropy, bandwidth, starting peak frequency and maximum peak frequency. PC3 loaded positively on entropy and starting peak frequency, and moderately negatively on average peak frequency and maximum peak frequency. Within three-dimensional PC space, the breeding songs of all four species occupied distinct regions largely separate from each other (Figure 2A, Supplementary Video, sample sizes below).

**Table 1:** Results of principal components analysis on the correlation matrix of nine acoustic parameters measured from *Prinia* acoustic signals.

|  | PC1 | PC2 | PC3 | PC4 | PC5 | PC6 | PC7 | PC8 | PC9 |
| --- | --- | --- | --- | --- | --- | --- | --- | --- | --- |
| **Average entropy** | -0.22 | 0.35 | 0.56 | -0.04 | 0.66 | -0.22 | -0.10 | -0.10 | -0.08 |
| **Bandwidth** | -0.37 | 0.36 | -0.21 | -0.02 | 0.09 | 0.50 | -0.21 | 0.09 | 0.61 |
| **Center frequency** | 0.41 | 0.26 | -0.15 | -0.19 | 0.14 | -0.23 | -0.14 | 0.78 | 0.04 |
| **Duration** | -0.41 | 0.06 | -0.21 | 0.04 | 0.03 | -0.42 | 0.73 | 0.15 | 0.21 |
| **Peak frequency** | 0.32 | 0.23 | -0.36 | -0.60 | 0.26 | 0.11 | 0.26 | -0.43 | -0.06 |
| **Peak frequency Start** | 0.24 | 0.44 | 0.49 | 0.05 | -0.36 | 0.37 | 0.47 | 0.10 | -0.05 |
| **Peak frequency End** | 0.33 | -0.07 | -0.23 | 0.66 | 0.52 | 0.27 | 0.23 | 0.00 | -0.04 |
| **Peak frequency Max** | -0.07 | 0.64 | -0.34 | 0.36 | -0.25 | -0.27 | -0.22 | -0.22 | -0.31 |
| **Peak frequency Min** | 0.44 | 0.03 | 0.17 | 0.16 | -0.11 | -0.40 | -0.07 | -0.32 | 0.69 |
| **Eigenvalue** | 4.32 | 1.67 | 0.98 | 0.64 | 0.59 | 0.29 | 0.28 | 0.13 | 0.09 |
| **% explained** | 47.97 | 18.54 | 10.87 | 7.15 | 6.58 | 3.23 | 3.18 | 1.46 | 1.01 |

**Figure 2:**
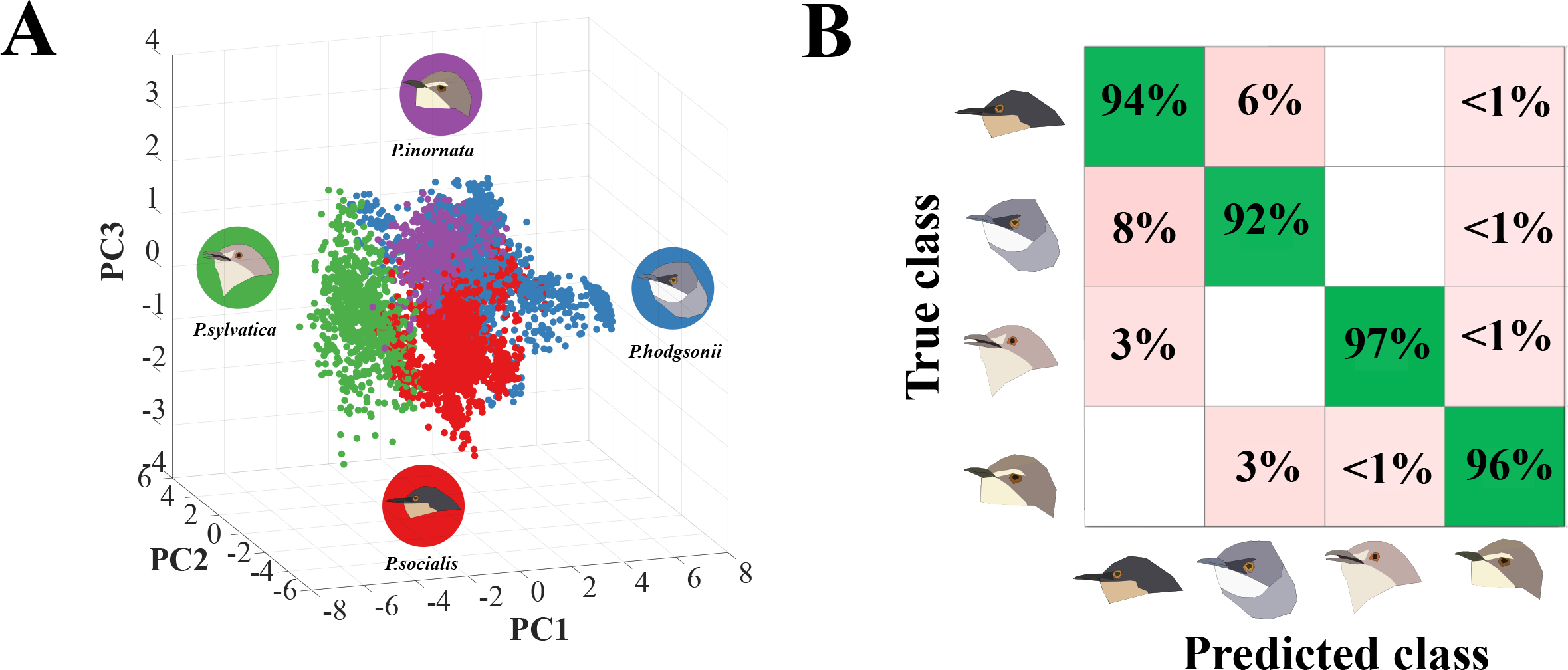
**A.** Principal component space (PC1-3) of the breeding songs of four *Prinia* species. Each species occupies a distinct region in this three-dimensional space. **B.** Confusion matrix of a linear discriminant classifier built in MATLAB using 9 acoustic parameters. Percentages in the green boxes indicate the correct classification rates for each species, and pink boxes the misclassification rates. Note the high degree of accuracy of LDA in assigning calls to species, indicating that they are divergent in signal space.

Linear Discriminant Analysis (LDA) further supported the observation of *Prinia* species occupying divergent regions of song parameter space from each other (Figure 2B). Our linear discriminant classifier could assign song notes to species with an overall accuracy of 94.2%. *P.socialis* (N=1799) had a correct classification rate of 94%, *P.hodgsonii* 92% (N=1356), *P.inornata* 96% (N=652) and *P.sylvatica* 97% (N=806). The most likely misclassification (only 6%) was of *P.socialis* notes being misclassified as *P.hodgsonii*, and an 8% probability of the converse. The high accuracy of the LDA model, and low misclassification rates further supports these four sympatric species occupying distinct regions of song signal space with little overlap. A similar LDA analysis on calls, classified separately from song notes, also uncovers some separation between species (Supplementary Figure 2). Although calls of *P.inornata* and *P.sylvatica* exhibited 99% and 100% correct classification rates in the LDA model, and *P.socialis* 86%, *P.hodgsonii* exhibited only a 55% correct classification rate, and was 45% likely to be misclassified as *P.socialis*. Thus, although *Prinia* are very distinct in song parameter space, the calls of at least two species exhibit similarity to each other.

Finally, the results of a randomization analysis in 9-dimensional PC space (Supplementary Figure 1, see Methods) also supported partitioning in song frequency-time parameters. Average interspecific distance was much greater (Z=63.04, P<0.001 using distributions fit to our original dataset, Z=7.386 * 10^12^, P<0.001 using randomly reassigned data points, see Methods) than expected by random chance (a distribution of 10000 randomized “null communities”). Additionally, each species pair (6 total pairs for 4 species) was more divergent in song than expected by random chance (Supplementary Data). Thus, our data are consistent with sympatric *Prinia* partitioning their breeding songs (but not necessarily call notes), and with each species occupying a unique region of 9-dimensional song parameter space.

### Interspecific differences in complexity of song repertoires

Partitioning of acoustic signal space exerts a constraint on intraspecific (within-species) signal diversity. We found that *Prinia* species exhibited different patterns in note diversity within intraspecific signal space. Using hierarchical clustering (see Methods), we identified 5 distinct note clusters in intraspecific principal component space for *P.socialis* (Figure 3A), and 7 for *P.hodgsonii* (Figure 3B). An LDA on these clusters further supports their distinctness from each other. The five clusters of notes in *P.socialis* all had correct classification rates between 81.3% and 100%, whereas the seven note clusters for *P.hodgsonii* had correct classification rates between 75.9% and 100% (Supplementary Figure 3). Preliminary examinations suggested that song bouts in *P.socialis* consisted of repetitions of a particular note type, whereas *P.hodgsonii* combined up to 4 note types in a single song bout (Supplementary Figure 4).

**Figure 3:**
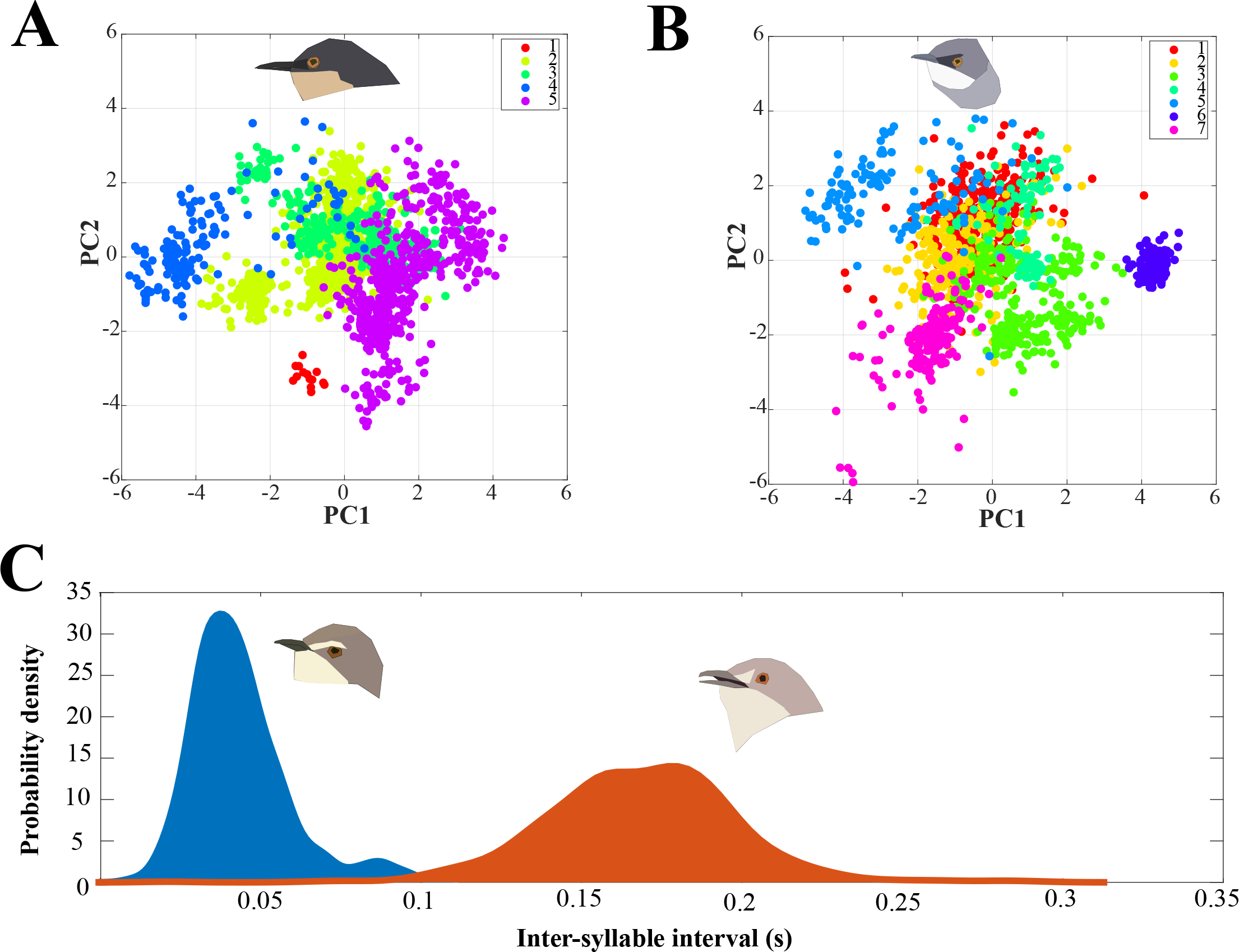
**A, B.** Note types of *P.socialis* (A) and *P.hodgsonii* (B). The different colors indicate distinct note types, recovered using hierarchical clustering on all 9 parameters and then mapped back onto two-dimensional PC space to visually depict the relative positions of each note type. **C.** Probability density plots of the time interval between successive notes for *P.inornata* (blue) and *P.sylvatica* (orange), two species with only one note type. Note that *P.inornata* sings a more rapid song (lower intervals), but that the peak is narrower. The broader peak of *P.sylvatica* suggests more intraspecific variation in repetition rate.

In contrast to the former two species, *P.inornata* and *P.sylvatica* both exhibited only a single note type with no clear clustering within intraspecific song parameter space (Supplementary Figure 3). We therefore investigated whether they exhibited intraspecific variation in the temporal pattern of their songs i.e. the inter-syllable interval. Probability density functions of the inter-syllable intervals revealed that the two species exhibit different strategies from each other (Figure 3C). *P.sylvatica* generally exhibited longer intervals but also a wider probability density distribution across all recorded vocalizations of this species, peaking between 150-200ms. This is consistent with the presence of faster and slower songs, and considerable intraspecific variation in syllable repetition rate. By contrast, *P.inornata* exhibited a narrow peak at approximately 40ms, indicating that its single note type is repeated rapidly at a more stereotyped rate, with relatively little variation in timing.

### Interspecific differences in song perch height

Finally, we examined interspecific patterns in song perch height to quantify whether *Prinia* species segregated from each other along this axis in acoustic signal space. Each species exhibited a distinct pattern in the use of song perches (Figure 4). *P. inornata* sang below 5m from the ground almost 60% of the time, and only very rarely from trees, whereas *P.sylvatica* sang almost entirely (>80% of the time) from trees, and only very rarely near the ground. *P.socialis* and *P.hodgsonii* sang mostly from intermediate heights, with the former distributing almost evenly from 2 to >10m, whereas the latter sang most commonly between 5-10m above the ground. Interspecific patterns in song perch heights were significantly different from chance (χ^2^ tests for categorical data: χ^2^= 34.03, dF=9, P<0.01, see Supplementary Data for more details). Thus, the species with the least complex repertoires (in terms of note diversity), generally sang from the highest and lowest song perches, whereas the two species with the greatest note diversity sang from intermediate heights within their scrub-grassland habitat.

**Figure 4:**
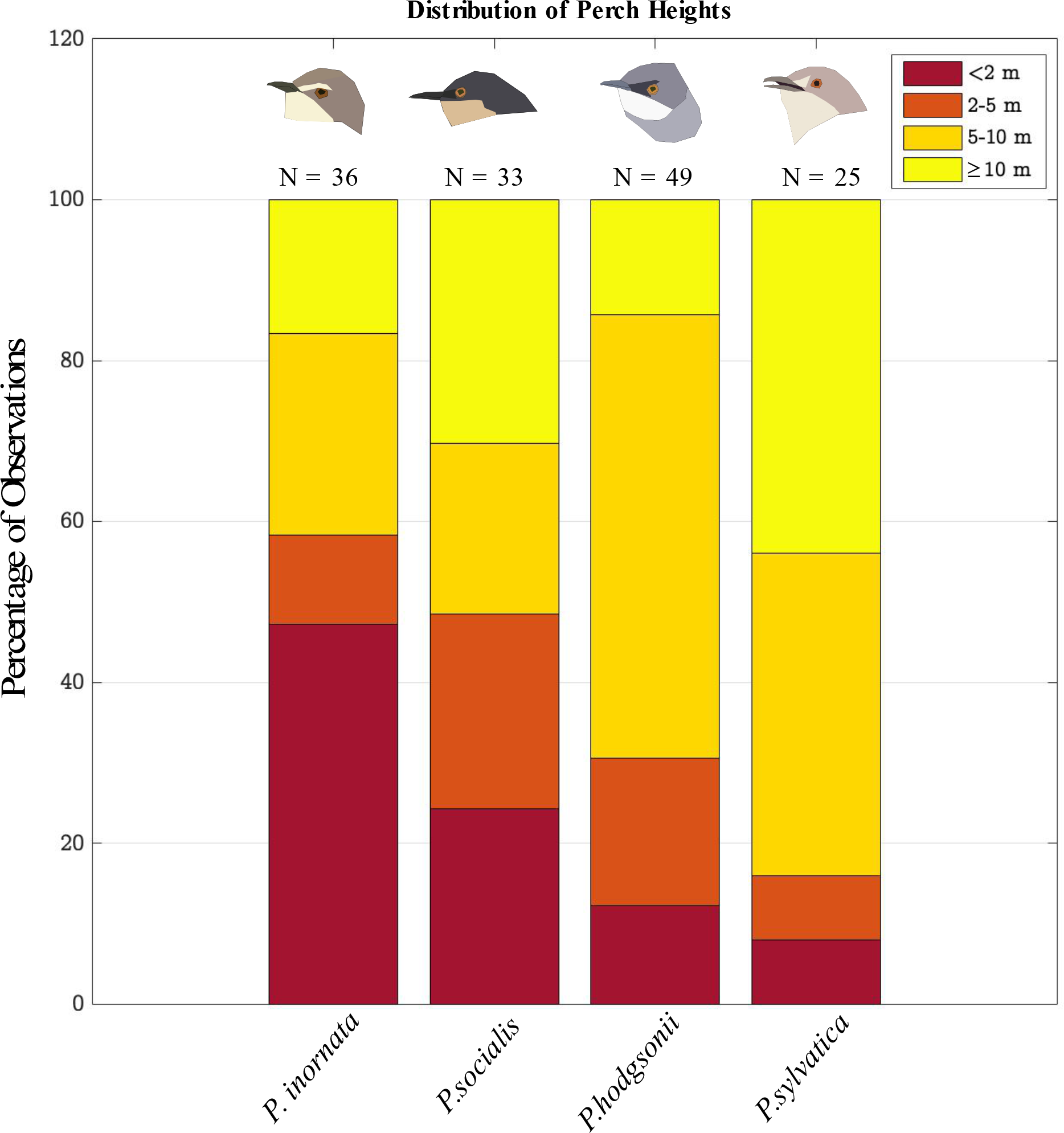
Stacked bar graphs depicting the distribution of song perch heights occupied by four *Prinia* species.

## Discussion

In summary, we uncover evidence that four sympatric passerine wren-warblers in a dry deciduous habitat in India are not only divergent in song parameters, but also exhibit partitioning in song perch height. This therefore results in a restricted intraspecific acoustic signal space along multiple axes, within which each species exhibits different note repertoires and diversity in communication signals. The two species that occupy the highest and lowest singing heights exhibit the most note stereotypy, and vary their note repetition rates to different extents. The two species occupying intermediate heights exhibit multiple different note types, which likely increases intraspecific signal variation within a constrained acoustic signal space. Below, we discuss how the patterns exhibited by these passerine birds are illustrative of the constraints that ecology and signal propagation may exert on signaling behavior.

### Acoustic signal space partitioning and the diversity of communication strategies

Partitioning of acoustic signal parameters minimizes overlap and masking interference(Schmidt et al. 2013), but additionally exerts a constraint on intraspecific signal diversity. Many organisms exhibit partitioning in frequency-time parameters of acoustic signals, including frogs, crickets and barbets(Chek et al. 2003; Schmidt et al. 2013; Krishnan and Tamma 2016). Both frequency and temporal traits are important facilitators in separating competing sound streams from each other. The breeding songs of *Prinia* are issued primarily during the monsoon breeding season(Rasmussen and Anderton 2005). In the landscape of seasonally dry scrub-grassland that characterizes large parts of Peninsular India, the four species studied here are some of the most abundant and vocal birds(Krishnan 2019), and are broadly sympatric over much of their range. Partitioning acoustic signal space is thus likely to be important in minimizing masking interference between them.

However, in addition to staying separate from heterospecifics, conspecific individuals must also modify their signals and signaling behavior in order to remain distinct from each other. A restricted acoustic signal space as a result of physical constraints and niche partitioning amplifies this problem. Our data suggests that wren-warblers overcome the constraints of their signal space on intraspecific signal diversity in different ways. *P.inornata* possesses only a single note type, and appears to sing a relatively stereotyped song without much variation. *P.sylvatica*, however, which also has a single note type, exhibits a broad range of repetition rates, consistent with possessing “faster” and “slower” songs. Finally, *P.hodgsonii* and *P.socialis* exhibit 7 and 5 note types respectively. These may correspond both to greater note diversity within a bout (as in *P.hodgsonii*), different song types (potentially what we observe in *P.socialis*) or to individual variation in notes, which requires further study involving banding birds. Our study examines note diversity only at the population level; what we do demonstrate, however, is that closely related bird species may evolve intraspecific signal diversity in multiple ways within the constraints of their acoustic niche.

Finally, *Prinia* species possess multiple call notes (including contact and alarm calls)(Rasmussen and Anderton 2005), in addition to their breeding songs. When examining interspecific differences in call notes, we find that at least two species (*P.socialis* and *P.hodgsonii*) may possess relatively similar call structures (unlike their songs which are very divergent), with high misclassification rates in an LDA analysis (Supplementary Figure 2). This may indicate that their calls exhibit some degree of convergence in acoustic structure, possibly owing to conserved function (for instance, alarm calls)(Bradbury and Vehrencamp 2011), and is a potentially interesting subject of future study.

### Perch height differences and the propagation of acoustic signals

In addition to partitioning their acoustic signals along frequency and time axes, a number of animal taxa are also known to segregate along a spatial axis by selecting different singing locations(Hodl 1977; Diwakar and Balakrishnan 2007). In many animals, this may help reduce interference effects, although other studies have pointed out that height differences may be less effective in reducing masking interference, especially if the overall height of available perches is low(Schmidt et al. 2013). Singing height is known to influence the distance of sound propagation(Marten and Marler 1977), and forest birds with lower pitches and slower songs sing closer to the ground where their signals propagate further(Nemeth et al. 2002). We find that the four *Prinia* species sing at different song perch heights. Interestingly, the two species with the fewest note types (*P. inornata* and *P.sylvatica*) sing at the lowest and highest heights, respectively, and *P. inornata*, which sings very close to the ground, has the faster-repeating song of the two. The other two species, which sing multiple note types (*P. socialis* and *P.hodgsonii*) occupy intermediate perch heights.

*P.sylvatica*, which consistently sings at the highest perches, also has the lowest average peak frequency and longest duration of its song notes, but the highest bandwidth on average of any of the four species (Table 2). This runs contrary to studies of birds from closed rainforest, where lower-pitched, slower songs are found closer to the forest floor(Nemeth et al. 2002). One possible reason for this is that our study was conducted in relatively open grassy habitats as opposed to the closed-canopy forests where other such studies were undertaken. It is possible that in such conditions, the higher bandwidths and longer notes of *P.sylvatica* enable longer-distance communication. The narrow-band, rapid songs of *P.inornata* may enable it to avoid detection by predators when singing very close to the ground. Studies have demonstrated a link between song perch selection and predation risk(Krams 2001). Field experiments may help address these questions by calculating relative signal propagation distances at various heights. An alternate possibility is that perch height partitioning is driven by other ecological factors. *P.sylvatica* is the largest of the four species(Grimmett et al. 1998), and we have observed aggression towards other *Prinia* by this species, as well as by *P.socialis*, the next species in size. Perch height partitioning may simply result from competition for higher song perches in this open environment, with *P.inornata* being driven to lower perches (intraspecific competition has been observed in nightingales)(Sprau et al. 2012). In support of this, all four species sing at least occasionally from the highest perches (Figure 4). Our data overall are consistent with a multidimensional acoustic niche in scrub-grassland passerine birds, with partitioning along multiple axes including frequency-time parameter space and physical singing space. The physical axis of song perch height is putatively related to an ecological dimension, i.e. competition for song perches and its consequences on signal propagation. Thus, we hypothesize that perch height partitioning by itself may not play a role in segregating the acoustic signal space of wren-warblers. By studying four closely-related species from the same genus, we also highlight the diversity of communication strategies employed by singing birds to minimize masking overlap while simultaneously generating diverse and distinct communication signals.

**Table 2:**
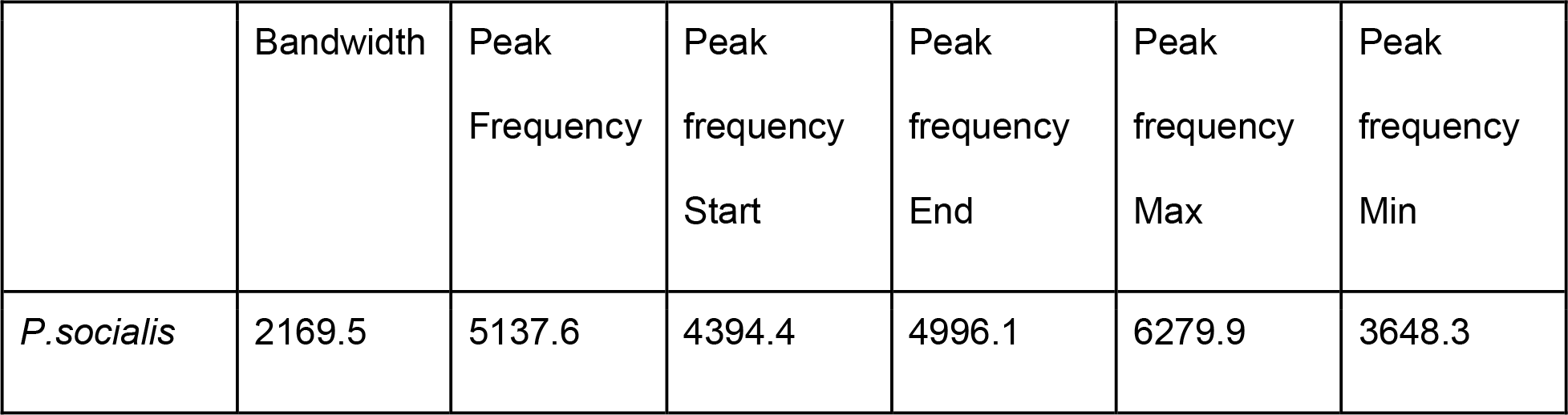
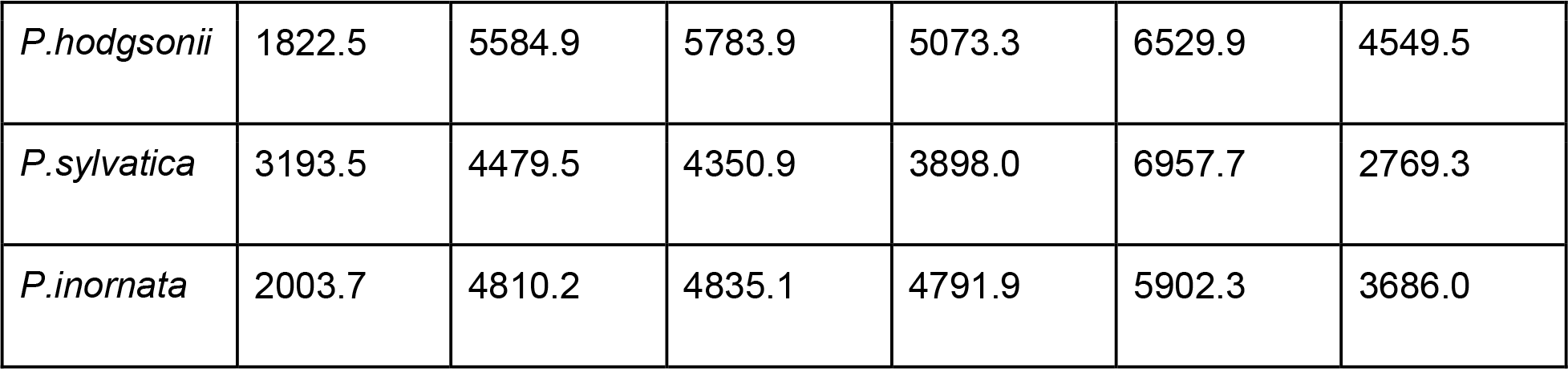
Frequency parameters and bandwidth (in Hz) of all four species. Values represent averages for each parameter for all measured notes.

## Acknowledgments

We thank Deepak Barua for help with equipment, Raghav Rajan and his lab for feedback and discussions, and Samira Agnihotri for comments on the data.

---

**Supplementary Figure 1:** q-q plots showing fits of each principal component score (PC1-9) to a normal distribution (straight line). Because the data along each PC approximately follow normal distributions, we fit random normal distributions to each and randomly resampled these to test for non-randomness of acoustic community structure. See Methods for more details. Bottom right: Frequency histogram of average interspecific distance for 10000 randomized “null” assemblages of species, using these randomly generated distributions. The observed interspecific distance in acoustic space is much larger, indicating partitioning of acoustic signal space.

**Supplementary Figure 2:** Confusion matrix indicating LDA misclassification rates for the calls of *Prinia* species. Color schemes are the same as in Figure 2.

**Supplementary Figure 3:** Top row: Song note PC space plots for *P.inornata* and *P.sylvatica*; all song notes fall into a single cluster. Bottom row: Confusion matrices indicating LDA misclassification rates for the note types of *P.socialis* (left) and *P.hodgsonii* (right).

**Supplementary Figure 4:** Spectrograms of different note types in *P.socialis* (top), and *P.hodgsonii* (bottom, white box). Note that song bouts in *P.socialis* consist of repetitions of a single note type, whereas *P.hodgsonii* incorporates multiple note types within a single song bout.

