## Supplementary figures and images for "Sympatric wren-warblers partition acoustic signal space and song perch height"

### Supplementary Figure 1

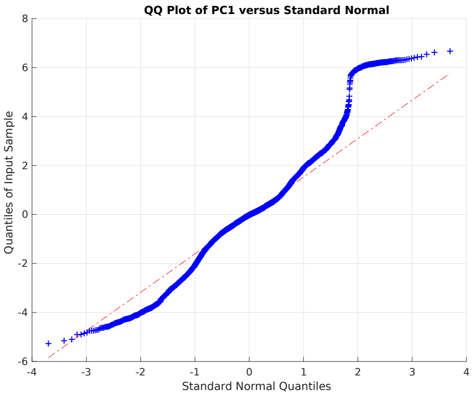


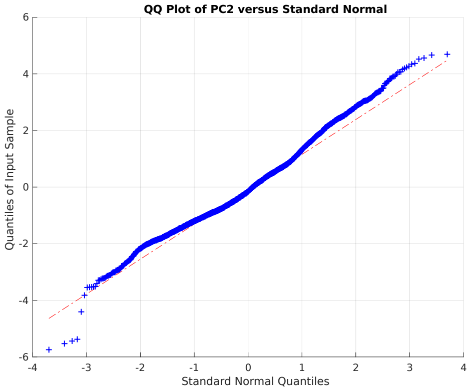


**PC1 (left) and PC2 (right)**

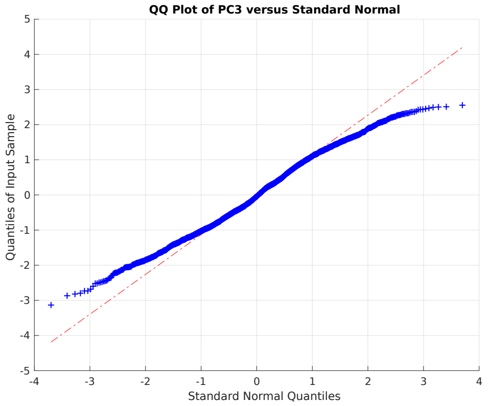


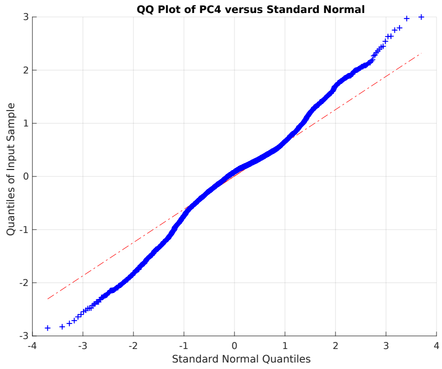


**PC3 (left) and PC4 (right)**

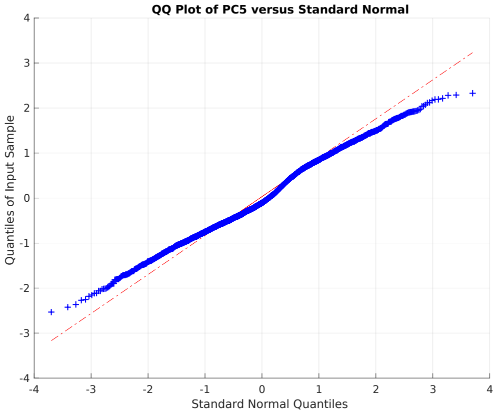


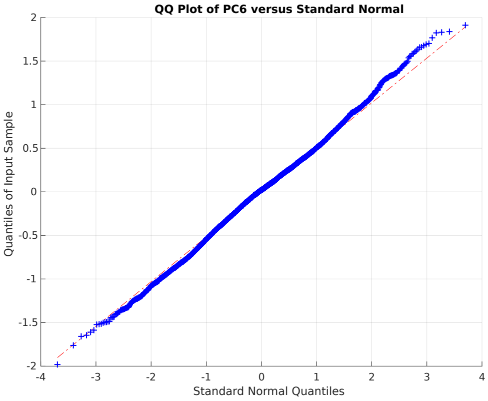


**PC5 (left) and PC6 (right)**

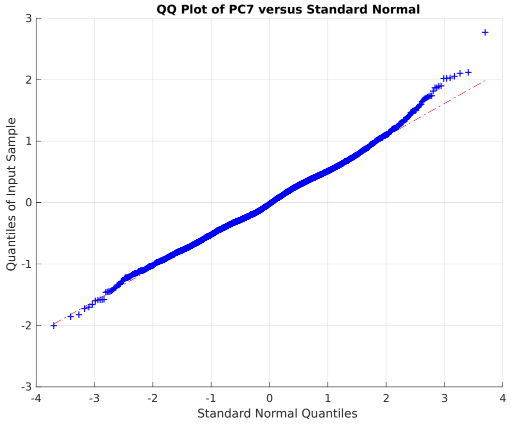


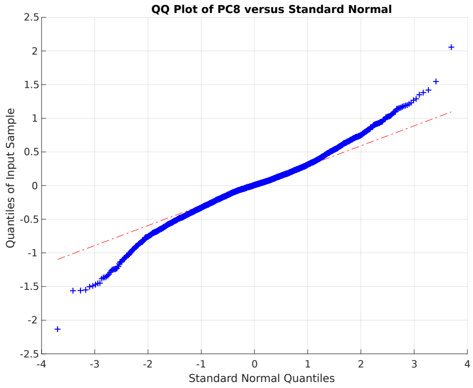


**PC7 (left) and PC8 (right)**

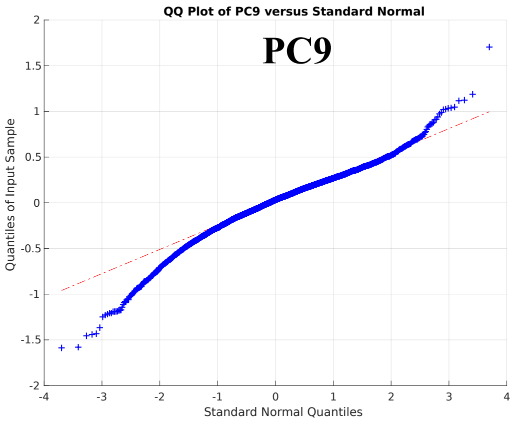


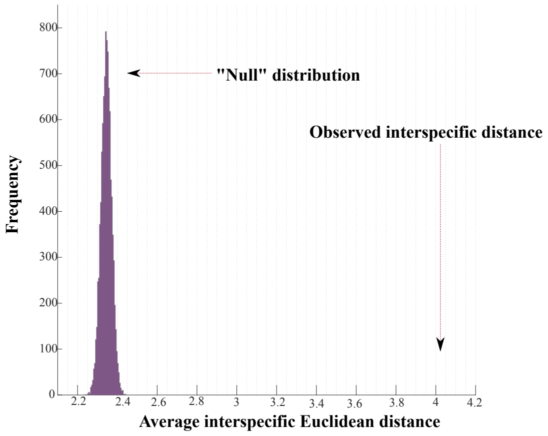

### Supplementary Figure 2

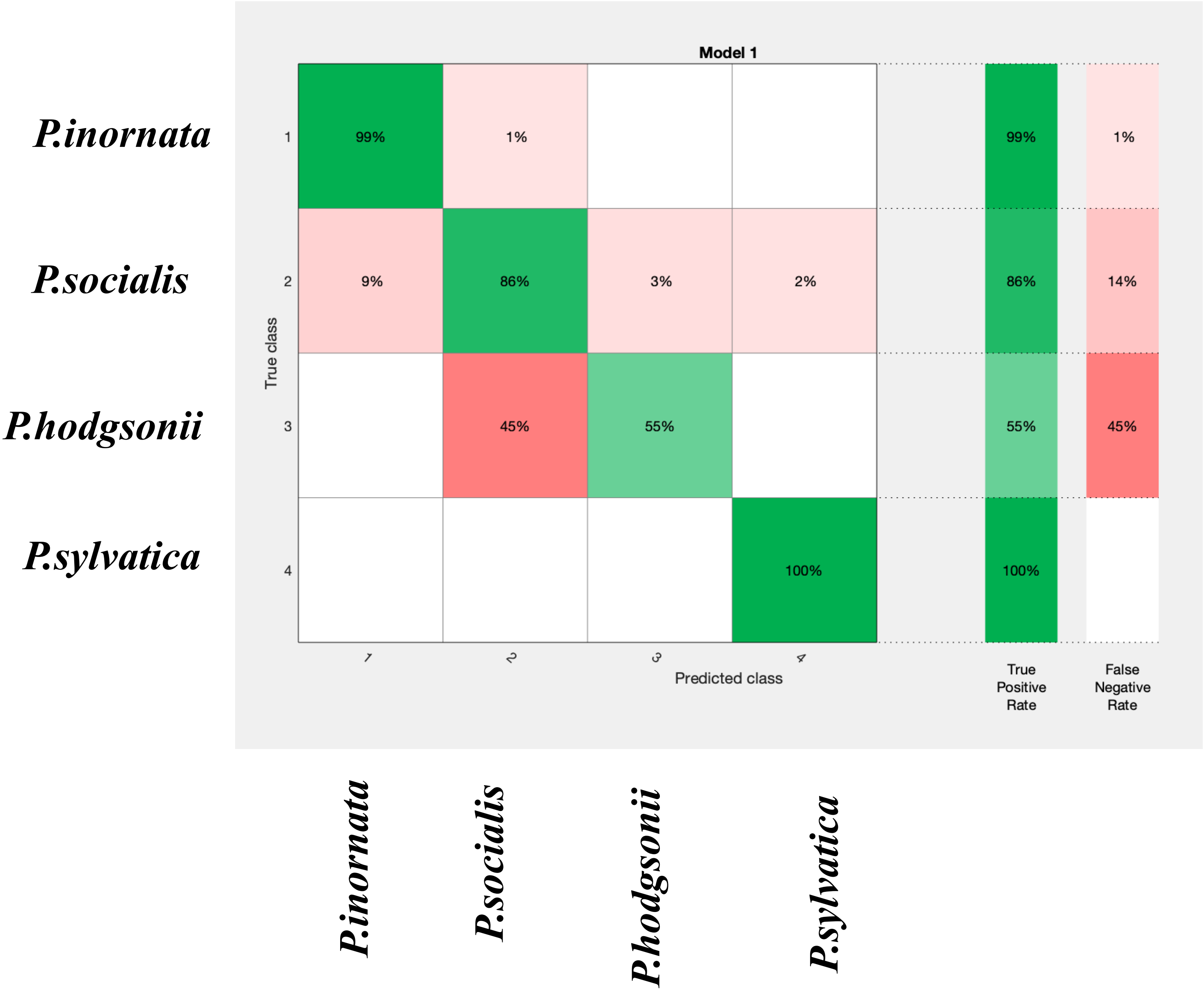

### Supplementary Figure 4

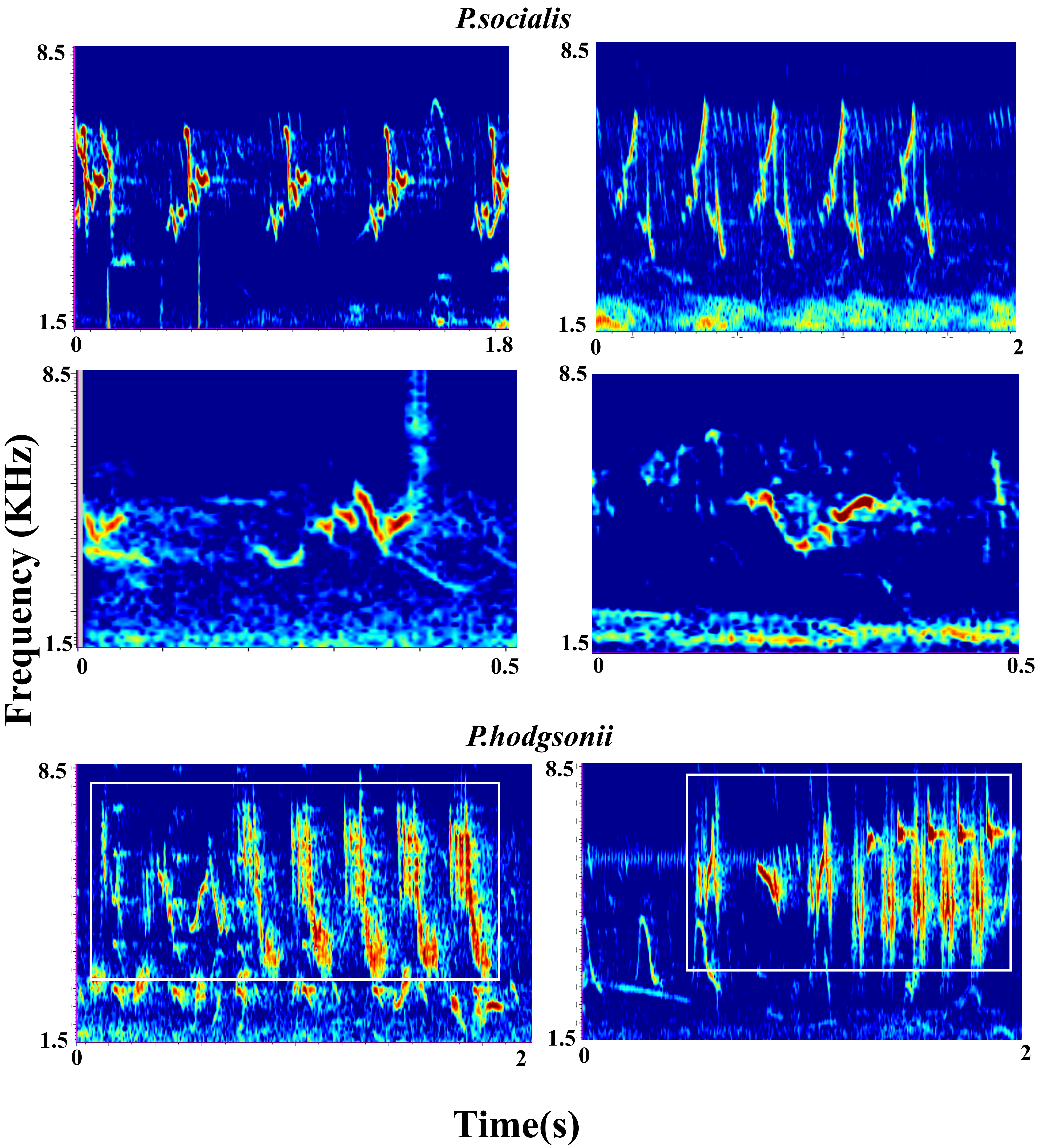
