## Supplementary Figure 3 for "Sympatric wren-warblers partition acoustic signal space and song perch height"

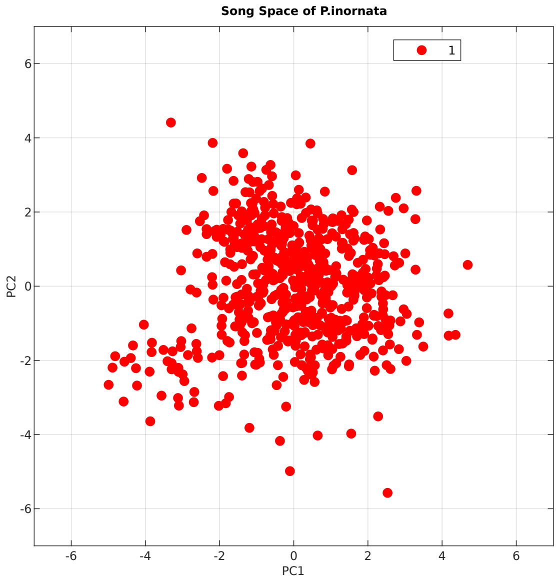


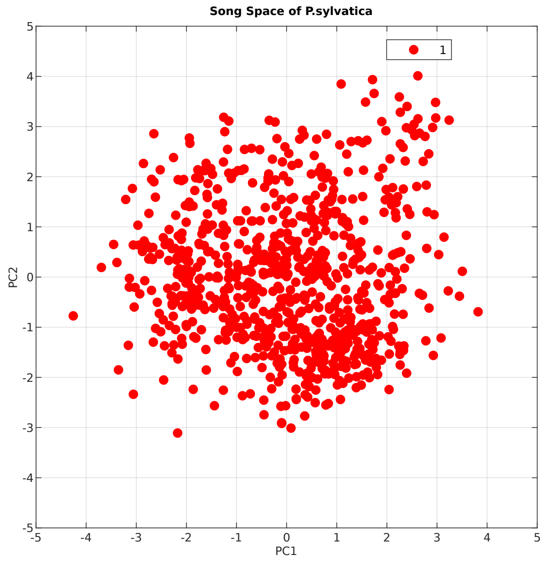


**Running LDA Classifier on classification based on clustering for Ashy Prinia**

Overall Accuracy of Model : 91.7%
Overall Error: 8.3%

Confusion Matrix

Rows: Classification based on clustering; Columns: Classes Predicted by LDA Classifier

| | 1 | 2 | 3 | 4 | 5 | TPR/FNR |
|---|---|---|---|---|---|---|
| 1 | 143<br>100% | | | | | 100%<br>0.0% |
| 2 | 1<br>0.1% | 712<br>97.1% | 4<br>0.5% | 9<br>1.2% | 7<br>1.0% | 97.1%<br>2.9% |
| 3 | | 16<br>5.3% | 274<br>90.7% | | 12<br>4.0% | 90.7%<br>9.3% |
| 4 | | 33<br>17.1% | 3<br>1.6% | 157<br>81.3% | | 81.3%<br>18.7% |
| 5 | 36<br>8.4% | 28<br>6.5% | | | 364<br>85.0% | 85.0%<br>15.0% |

**Running LDA Classifier on classification based on clustering for Grey Breasted Prinia**

Overall Accuracy of Model : 83.8%
Overall Error: 16.2%

Confusion Matrix

Rows: Classification based on clustering; Columns: Classes Predicted by LDA Classifier

| | 1 | 2 | 3 | 4 | 5 | 6 | 7 | TPR/FNR |
|---|---|---|---|---|---|---|---|---|
| 1 | 187<br>84.6% | 21<br>9.5% | 1<br>0.5% | 2<br>0.9% | 10<br>4.5% | | | 84.6%<br>15.4% |
| 2 | 29<br>8.6% | 257<br>76.3% | 3<br>0.9% | 8<br>2.4% | 15<br>4.5% | | 25<br>7.4% | 76.3%<br>23.7% |
| 3 | 2<br>0.8% | 22<br>9.1% | 201<br>83.1% | 11<br>4.5% | | 6<br>2.5% | | 83.1%<br>16.9% |
| 4 | 10<br>10.3% | | 1<br>1.0% | 86<br>88.7% | | | | 88.7%<br>11.3% |
| 5 | 19<br>13.9% | 14<br>10.2% | | | 104<br>75.9% | | | 75.9%<br>24.1% |
| 6 | | | | | | 148<br>100% | | 100%<br>0.0% |
| 7 | | 19<br>10.9% | 1<br>0.6% | | | | 154<br>88.5% | 88.5%<br>11.5% |
